# A population-genomic approach for estimating selection on polygenic traits in heterogeneous environments

**DOI:** 10.1101/2020.06.02.129874

**Authors:** Zachariah Gompert

**Affiliations:** Department of Biology, Utah State University, Logan, UT 84322, USA; Ecology Center, Utah State University, Logan, UT 84322, USA

**Keywords:** fluctuating selection, ecological genetics, polygenic traits, approximate Bayesian computation, computational statistics, *Callosobruchus maculatus*

## Abstract

Strong selection can cause rapid evolutionary change, but temporal fluctuations in the form, direction and intensity of selection can limit net evolutionary change over longer time periods. Fluctuating selection could affect molecular diversity levels and the evolution of plasticity and ecological specialization. Nonetheless, this phenomenon remains understudied, in part because of analytical limitations and the general difficulty of detecting selection that does not occur in a consistent manner. Herein, I fill this analytical gap by presenting an approximate Bayesian computation (ABC) method to detect and quantify fluctuating selection on polygenic traits from population-genomic time-series data. I propose a model for environment-dependent phenotypic selection. The evolutionary genetic consequences of selection are then modeled based on a genotype-phenotype map. Using simulations, I show that the proposed method generates accurate and precise estimates of selection when the generative model for the data is similar to the model assumed by the method. Performance of the method when applied to an evolve-and-resequence study of host adaptation in the cowpea seed beetle (*Callosobruchus maculatus*) was more idiosyncratic and depended on specific analytical choices. Despite some limitations, these results suggest the proposed method provides a powerful approach to connect causes of (variable) selection to traits and genome-wide patterns of evolution. Documentation and open source computer software (`fsabc`) implementing this method are available from GitHub (https://github.com/zgompert/fsabc.git).

## Introduction

Selection can cause rapid evolutionary change on ecological time scales (Ford, 1977; Reznick *et al*., 1997; Bell, 2010; Thompson, 2013; Messer *et al*., 2016). Rates of evolution are lower when measured on longer time scales (Hendry & Kinnison, 1999; Kinnison & Hendry, 2001; Gingerich, 2019), in part because the form, direction and intensity of selection often vary through time (e.g., from generation to generation; Siepielski *et al*., 2009; Bell, 2010). Various factors can cause selection to fluctuate in time, including climatic variability (Grant & Grant, 2002; Bergland *et al*., 2014; Grant & Grant, 2014; Siepielski *et al*., 2017), host-parasite or host-pathogen coevolution (Gómez & Buckling, 2011; Hall *et al*., 2011; Ford *et al*., 2017), and mating or survival advantages for rare phenotypes (Endler, 1988; Takahashi *et al*., 2010; Hughes *et al*., 2013; Nosil *et al*., 2018). Similar factors can cause selection to vary in space (Kingsolver *et al*., 2001; Siepielski *et al*., 2013), resulting in local adaptation (Endler, 1977). Fluctuating selection may help explain fundamental biological phenomena, such as the maintenance of polymorphism (Ford, 1977; Turelli *et al*., 2001; Hedrick, 2006; Wittmann *et al*., 2017), molecular diversity levels (Gillespie, 1991; Hahn, 2008; Leffler *et al*., 2012; Coop, 2016), ecological specialization (Agrawal *et al*., 2010; Anderson *et al*., 2013; Gompert & Messina, 2016; Agrawal, 2020) and the evolution of plasticity (Hallsson & Björklund, 2012; Tufto, 2015; King & Hadfield, 2019). Quantitative studies of the population genetic consequences of fluctuating selection in the wild are necessary to better evaluate this possibility, but remain relatively rare (Messer *et al*., 2016) (but see, e.g., Ford, 1977; Mueller *et al*., 1985; Bergland *et al*., 2014).

Such studies have been hampered, in part, by limited development of appropriate statistical methods (Messer *et al*., 2016). Many popular statistical methods infer selection on genetic variants from static population-genomic patterns (i.e., patterns lacking a temporal component), such as haplotype structure and diversity (Sabeti *et al*., 2002; Szpiech & Hernandez, 2014), genetic differentiation between populations (Beaumont & Nichols, 1996; Foll & Gaggiotti, 2008; Nosil *et al*., 2008), or genotype–environment associations (Coop *et al*., 2010; De Villemereuil & Gaggiotti, 2015; Rellstab *et al*., 2015). These methods can detect selection when it occurs in a consistent manner over time, but were not designed to identify temporally fluctuating selection. Methods (e.g., Illingworth & Mustonen, 2011; Mathieson & McVean, 2013; Feder *et al*., 2014; Gompert, 2016; Buffalo & Coop, 2019; Kelly & Hughes, 2019), and associated data sets (e.g., Bi *et al*., 2013; Reich, 2018; Rêgo *et al*., 2019; Bi *et al*., 2019), with temporal sampling in natural or experimental populations are better suited for detecting fluctuating selection and are becoming more common.

Mueller *et al*. (1985) proposed one of the first explicit tests for fluctuating selection, which was based on the proposition that genetic loci evolving in response to the same (unknown) environmental variations should exhibit correlated patterns of allele frequency change over time. Using this method, Mueller *et al*. (1985) found ample evidence of fluctuating selection on allozyme loci in natural populations of two *Drosophila* species. More recently, Bergland *et al*. (2014) found evidence of genomically-widespread fluctuating selection in *D. melanogaster* based on repeated seasonal oscillations in the frequency of multiple SNP markers. Gompert (2016) introduced a method to quantify fluctuating selection on genetic loci from correlations between patterns of allele frequency change and the state of the environment. This requires an explicit hypothesis about the environmental factor causing selection to fluctuate. Buffalo & Coop (2019) derived a method to quantify the extent and consequences of (linked) selection from population-genomic time series from autocovariance in allele frequency changes across generations. This method can be used to detect fluctuating selection, provided shifts in the direction of selection do not occur frequently (Buffalo & Coop, 2019).

With the exception of Buffalo & Coop (2019), methods designed to detect fluctuating selection perform best when individual genetic loci experience strong selection. However, many traits are polygenic (e.g., Pritchard *et al*., 2010; Yang *et al*., 2010; Shi *et al*., 2016; Lucas *et al*., 2018; Gompert *et al*., 2019), and thus, selection on genes can be weak even when selection on a trait is intense (Walsh & Lynch, 2018). A general approach to overcome this limitation was suggested by Berg & Coop (2014) (also see Josephs *et al*., 2019). Specifically, polygenic adaptation can be inferred by incorporating genotype-phenotype associations from genome-wide association studies (GWAS) in population-genomic tests for selection. This makes it possible to accumulate evidence of selection across trait-associated loci, and thus detect selection even when none of the individual loci experience strong selection. Thus far, this analytical framework has mostly been applied to static population-genomic data sets, and not to detect fluctuating selection from temporal data.

Herein, I fill this analytical gap by presenting an approximate Bayesian computation (ABC) method to detect and quantify fluctuating selection on polygenic traits from timeseries data. With this method, phenotypic selection is modeled as an explicit function of the state of the environment (similar to Gompert, 2016). The population-genomic consequences of selection are then modeled based on estimated genotype-phenotype associations (similar to Berg & Coop, 2014). This allows inferences to be informed by patterns of change across multiple genetic loci, populations, and generations. Pooling information in this way increases statistical power, but precludes generic genome scans for selection. Although this could be viewed as a limitation, by making the genotype-phenotype-environment-fitness hypothesis explicit, the proposed approach places an emphasis on the ecological interactions that cause selection to fluctuate and thereby provides more meaningful inferences about the evolutionary process (as advocated for in a general sense by Endler, 1986).

In this study, I first describe the theoretical basis for the approach. This is followed by a description of the proposed ABC method, including the data required and a framework for model evaluation and comparison. The efficacy and limitations of the proposed method are then evaluated by applying it to thousands of simulated data sets, and to a recent experimental evolution study of host adaptation in seed beetles (*Callosobruchus maculatus*). I show that the proposed method produces precise estimates of selection, especially when the generative model for the data coincides with the model assumed by the method, and detects selection on weight in *C. maculatus* despite the low heritability of this trait. I conclude by discussing the utility and limitations of the proposed method, along with potential extensions of the proposed method.

## Methods

### Model

I first describe a mathematical model that connects phenotypic selection on a quantitative trait to allele frequency change at causal or linked loci (that is, loci affecting the trait or those in linkage disequilibrium with such loci). Selection is allowed to fluctuate in space and time because of variation in the environment. Selection on the trait is connected to expected allele frequency change based on an understanding of the selected trait’s genetic architecture, as might be obtained from a genome-wide association mapping study. If causal loci and their phenotypic effects are known, the selection estimated is the direct selection on the causal variants, otherwise it is the indirect selection arising from linkage disequilibrium (Lande & Arnold, 1983; Gompert *et al*., 2017). The goal of the model is to estimate the intensity of selection on the trait and how this varies by environment, as well as the extent to which this translates into selection on different genetic loci. Key assumptions of the model are that selection is always directional, that each locus explains a small proportion of the phenotypic variance (either because the trait is highly polygenic or has a large environmental variance), that the phenotypic effects of loci do not depend on the environment and that the causal variants (or linked loci) exhibit minimal linkage disequilibrium with each other such that each experiences genetic drift independently (I reiterate and expand on these assumptions below). I also assume that, at least over the duration of the population-genomic time series, mutation and gene flow are negligible and can be ignored. After introducing the model in this section, I describe the proposed approximate Bayesian computation method for fitting the model in the next section.

Let *S*_*jk*_ = *µ ∗ –µ* denote the selection differential for a quantitative trait in population *j* and generation *k*. Here, *µ* and *µ∗* denote the trait mean before and after selection, respectively. I assume phenotypic selection is a function of the environment. I consider three alternative models for the relationship between that state of the environment, denoted *x*_*jk*_, and the selection differential, *S*_*jk*_, specifically:

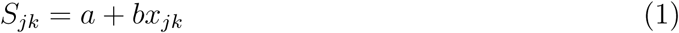

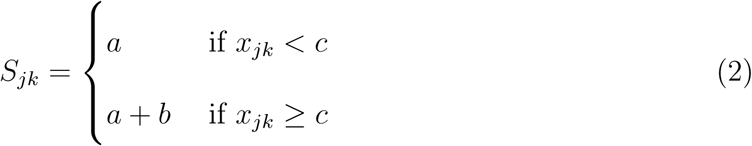

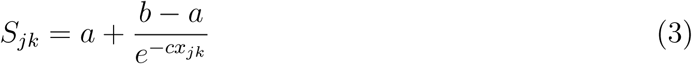

Here, *a, b* and *c* are coefficients that determine the effect of the environment on the selection differential; equations 1, 2 and 3 specify a linear, step and sigmoidal relationship between the environment and selection differential, respectively (Fig. 1A). The model thus assumes that phenotypic selection is always directional and independent of the current mean trait value, but allows flexibility in how the environment affects the direction and intensity of selection. Furthermore, I focus on the simplest case where selection is a function of a single environmental variable, but the extension to multiple environmental variables is trivial.

**Figure 1:**
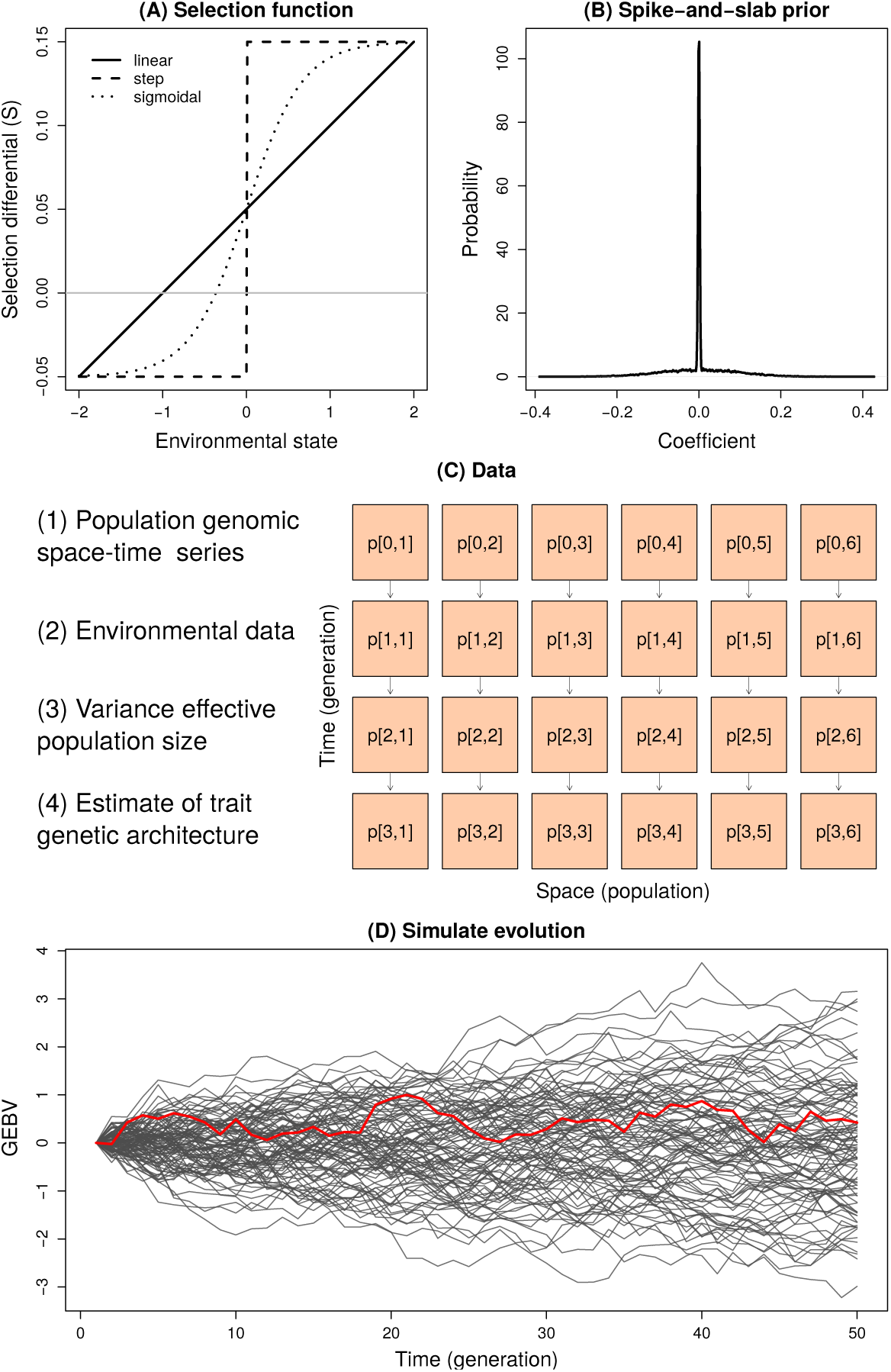
Graphical overview of the model, data and proposed method. Panel (A) shows hypothetical linear (Eqn. 1), step (Eqn. 2) and sigmoidal (Eqn. 3) functions for the effect of the environment on the selection differential. Panel (B) depicts the general form of the spike-and-slab priors used for the coefficients in the selection functions. Most of the prior probability is near zero, but large values are not improbable. Panel (C) summarizes the data for the approximate Bayesian computation (ABC) method. The four key data inputs are listed along with a diagram summarizing the population genomic time series, which includes allele frequency data (denoted *P*) for multiple loci measured across populations (space) and generations (time). The ABC approach simulates evolutionary trajectories for genetic loci and genomic-estimated breeding values (GEBVs) for each population based on the input data from (C). Panel (D) shown a hypothetical observed GEBV trajectory (red line) and additional simulated trajectories (gray lines) with different values for selection. The ABC approach retains the set of simulations that produce outputs most similar to the observed data (see the main text for details).

I then specify a model that relates phenotypic selection to expected allele frequency change. Let *w*_*i*_ denote the marginal, relative fitness of the non-reference allele at locus *i* (bi-allelic loci are assumed), that is *w*_*ijk*_ = ∫_*z*_*w*_*jk*_(*z*)*f*_*i*_(*z*)*dz*. Here, *w*_*jk*_(*z*) is the expected fitness of an individual with trait value *z* and *f*_*i*_(*z*) is the phenotypic distribution (density function) for individual’s carrying a copy of the non-reference allele at locus *i* (this includes heterozygotes and homozygotes for the non-reference allele) (note, at present I assume that *f*_*i*_(*z*) is fixed, that is I assume no plasticity). Following Kimura & Crow (1978) and Walsh & Lynch (2018), the intensity of selection on the non-reference allele at locus *i* can be approximated as

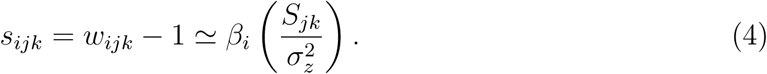

Here, *β*_*i*_ denotes the average phenotypic excess of the non-reference allele at locus *i*, which is defined in terms of mean trait values and frequencies of the three genotypes, 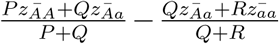, where *P*, 2*Q* and *R* are the frequencies of homozygous non-reference, heterozygous and homozygous reference genotypes and 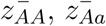 and 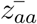 are their respective mean trait values (Fisher, 1941). If the genotypes at each causal locus are in their Hardy-Weinberg expected proportions and in linkage equilibrium across loci, this is equivalent to the average effect of an allele, as estimated with standard genome-wide association mapping methods (e.g., Zhou & Stephens, 2012; Zhou *et al*., 2013). The term *s*^2^ in Eqn. 4 denotes the phenotypic variation, and thus 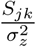 is the standardized selection differential (i.e., the selection differential in units of standard deviations). Thus, the selection on an individual allele is approximately equal to the product of the intensity of phenotypic selection and average phenotypic excess of the allele (Walsh & Lynch, 2018). This approximation is true or approximately true for a variety of fitness functions, but requires that phenotypic variation caused by any one allele is small relative to the phenotypic variance, and thus that selection on each individual allele is weak. This assumption can hold even when phenotypic selection is strong, if the trait is polygenic and lacks common, major effect loci.

Importantly, *s*_*ijk*_ measures selection as the average excess in relative fitness for the non-reference allele at locus *i*, not the more commonly used selection coefficient (which defines selection relative to a specific genotype, e.g., Gillespie, 2004). The average excess in relative fitness directly measures the effect of selection in bringing about allele frequency change, as such, this metric is not static but depends on the allele frequencies (Kimura & Crow, 1978; Walsh & Lynch, 2018). The expected change for the non-reference allele frequency at locus *i* due to selection is given by Δ*p*_*ijk*_ = *p*_*ijk*_*s*_*ijk*_. Genetic drift is assumed to also contribute to evolutionary change, such that,

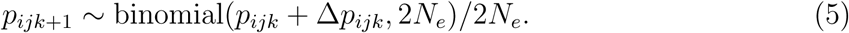

Equation 5 is the Wright-Fisher model with selection; *N*_*e*_ denotes the variance effective population size for population *j* and generation *k* (I omit subscripts on *N*_*e*_ for readability) (Ewens, 2004).

I assume that, conditional on the *s*_*ijk*_, evolutionary change is independent across genetic loci (i.e., linkage disequilibrium among the genetic loci is minimal). The validity of this approximation depends on the trait genetic architecture and set of genetic loci included in the analysis. For example, an assumption of linkage equilibrium is more reasonable when causal variants are dispersed across the genome rather than clustered in one or a few genomic regions. The approximation will also be better when each causal genetic variant is represented by a single genetic marker locus, rather than a set of linked genetic markers statistically associated with the same causal variant (see the “ABC approach” below for further discussion).

### ABC approach

The proposed method for estimating selection requires four data sources. First, estimates of allele frequencies in one or more populations for multiple time points (ideally consecutive generations) are required (these are assumed to be known with little or no error, but this assumption can be relaxed). Whole genome sequences are not needed (or warranted), but rather the method requires a set of genetic markers (e.g., SNPs) known to affect the trait of interest, or as will be more commonly the case, a set of markers statistically associated with the trait (i.e., in LD with the unknown causal variants). Second, information on the effects (associations) of the genetic markers with the trait are needed. Here, I assume that each marker has a probability of affecting or being associated with the trait, denoted *γ*_*i*_ (e.g., a posterior inclusion probability or PIP), and an effect conditional on a true association, denoted *β*_*i*_ (which I equate with *β*_*i*_ from Eqn. 4 assuming the genotypes in the populations are approximately equal to their expected, Hardy-Weinberg proportions; see Fisher, 1941). Standard models for polygenic GWA mapping and genomic prediction, such as the Bayesian sparse linear mixed model approach from gemma, output this information (Zhou *et al*., 2013). Such models provide probabilistic estimates of direct phenotypic effects (and thus of direct selection with the model) when the causal variants are sequenced, but only indirectly approximate direct selection when this is not the case (Gompert *et al*., 2017). Alternatively, *γ* (the probability of association) can be set to 1 for a set of genetic variants to indicate they are known to directly affect the trait; such confidence might be appropriate if genotype-phenotype associations have been validated by genetic manipulations (e.g., Barrett *et al*., 2019). Lastly, model-averaged effect estimates can be used, that is 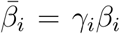, with the probability of association then set to 1. I explore this final possibility with an empirical data set.

Third, estimates of the variance effective population size for each population are needed. These can be estimated using either LD-based methods or change over time for a set of genetic markers not associated with the (putatively) selected trait (e.g., Jorde & Ryman, 2007; Do *et al*., 2014; Gompert & Messina, 2016). The proposed method accounts for uncertainty in estimates of *N*_*e*_ by integrating over a posterior distribution.

Fourth, the proposed method requires measurement of an environmental covariate hypothesized to determine the phenotype-fitness relationship (i.e., the selection differential). This covariate should be measured in each population and generation (or time step). I use the term environmental covariate in a broad sense, as this could include a variety of factors: temperature, precipitation, composite climatic variables from an ordination (e.g., principal components), predator abundance, resource availability, or even population density or the frequency of a trait in the population. Moreover, the relevant covariate value at time *t* could represent the measurement at time *t* or a cumulative measure over some past amount of time.

Given these data, I propose an approximate Bayesian computation (ABC) method for estimating the parameters of the selection function, *a, b* and *c* in Eqns. 1, 2 and 3, as well as derived parameters related to selection on the genetic loci, which I describe below. This method is implemented in the C++ program `fsabc` (version 0.1; available from GitHub), which is used in combination with the R package abc (version 2.1; Csillery *et al*., 2012) for inference. As with other ABC methods, the proposed method involves first sampling parameter values from prior probability distributions, then simulating evolution based on those parameter values, and finally calculating summary statistics for the simulated and observed data (all done with `fsabc`; Fig. 1). Posterior distributions for the parameters are then obtained based on the subset of sampled parameter values and simulations that most closely match the observed data in terms of the summary statistics (e.g., Sisson *et al*., 2018).

I assume the form of the selection function (linear, step or sigmoidal) is known, and that the coefficients describing the function (*a, b* and *c*) are the primary, unknown parameters (but see below for model selection and the possibility of model-averaging) (Fig. 1A). I specify spike-and-slab priors for each of these parameters, such that Pr(*a*) *∼ πδ*_0_ +(1*–π*)U(*l, u*) (Fig. 1B). Here, *π* is the probability of a zero coefficient and U(*l, u*) denotes a uniform probability density function with lower bound *l* and upper bound *u. δ*_0_ denotes a point mass at zero. This prior is conservative as it places a substantial proportion of the prior probability on zero (the exact value is given by *π*), but still allows for large coefficients (based on *l* and *u*). Values of *N*_*e*_ and the specific set of genetic markers associated with the trait (if either are expressed with uncertainty) are then sampled according to their probabilities.

Derived parameters *S*_*jk*_ (the selection differential) and *s*_*ijk*_ (selection on each locus) for population *j* and generation *k* are then calculated from Eqns. 1, 2 or 3, and Eqn. 4 based on the sampled parameter values. The simulation for each population is initiated with the known, initial allele frequencies in generation *k* = 0, and the allele frequencies in the following generation are then determined based on the stochastic Wright-Fisher model (Eqn. 5) (this is then the starting point for the next generation). This process is then repeated for each population and generation to simulate the complete population-genomic time series.

I compute two summary statistics for inference of the selection function parameters (*a, b* and *c*): (i) the total change in the mean polygenic score (i.e., mean genomic-estimated breeding value or GEBV), and (ii) the covariance between change in the mean polygenic score and the value of the environmental covariate. The total change in the mean polygenic score captures the average selection differential (e.g., *a* in the linear model, Eqn. 1), and is defined as Δ_*BV*_ = S_*ijk*_*β*_*i*_(*p*_*ijk*+1_ *– p*_*ijk*_). The covariance between change in the breeding value and the environmental state, that is COV(Δ_*BV*_*jk, x*_*jk*_), captures the manner in which the environment affects the selection differential (e.g., *b* in the linear model, Eqn. 1).

Primary interest is in inference of the selection function parameters, that is the parameters that describe how the environment affects the selection differential on the trait. However, the proposed method can also estimate derived parameters describing selection on each genetic locus, that is *s*_*ijk*_. I focus on two derived parameters that summarize generation, population and locus selection coefficients, specifically, the average absolute intensity of selection on the genetic loci (i.e., mean(|*s*_*ijk*_|)) and the standard deviation of the absolute intensity of selection (i.e., SD(|*s*_*ijk*_|)). These provide high-level summaries of the intensity and variability of selection on loci associated with or affecting the selected trait (i.e., indirect or direct selection depending on the nature of the genotype-phenotype map).

Various approaches have been developed to generate samples from an approximate posterior distribution in an ABC framework from summary statistics (Beaumont, 2010; Sisson *et al*., 2018). Here, I used the rejection with local-linear regression correction algorithm proposed by Beaumont *et al*. (2002) and implemented in the R abc package (Csillery *et al*., 2012). This approach performs quite well (see Results), but other approaches, such as ridge regression or non-linear regression (Blum & Francois, 2010) could be used instead for this portion of the analysis.

Beyond parameter estimation, the method also provides a convenient way to compute a posterior predictive distribution, which can be used for model assessment and comparison. For this, I simulate additional evolutionary trajectories by sampling parameter values (*a, b* and *c*) from their posterior distributions based on an initial model-fitting analysis (as described above). I then compute a new set of summary statistics for each simulated data set. Specifically, percentiles (10th to 90th percentiles in steps of 10%) of the distribution of change in the mean polygenic score across populations and generations are computed. I chose this novel set of summary statistics so that different information could be used for model fitting and model validation or comparison. By simulating multiple new data sets, posterior predictive distributions for each of the percentile summary statistics can be obtained and compared to the same summary statistics calculated from the observed data, with higher correspondence being indicative of better model performance.

### Analysis of simulated data

I first evaluated the performance of the method with simulated data. I began by testing how the method performed under a variety of conditions, but where the data were simulated under the same model used to analyze the data. This allowed me to assess the ability of the approach to provide precise and accurate parameter estimates when the model was the correct model for the data. I then conducted additional simulations to assess how the method performed for data simulated under alternative models.

The first set of simulations was conducted using the model described above and with the same software used for inference (i.e., `fsabc`). Standard (baseline) conditions involved 10 populations sampled in 10 successive generations. I further assumed 100 genetic loci affected the focal trait. Initial allele frequencies for these loci were sampled from a beta distribution with shape parameters equal to 0.5. The phenotypic effects of these 100 loci were drawn from a centered normal distribution. The standard deviation was set to ensure a heritability of *∼*0.3 (with a phenotypic variance of 1.0). That is, I assumed the trait was polygenic, lacked major effect loci, and had a modest heritability (and thus a substantial non-genetic variance component). I assumed the variance effective population size was estimated with uncertainty, such that knowledge and uncertainty in *N*_*e*_ was characterized by a re-scaled beta distribution, *N*_*e*_ *∼* beta(*α* = 10, *β* = 10, min = 400, max = 600) (here min and max are the lower and upper bounds). This distribution has a mean of *N*_*e*_ = 500 and standard deviation of 21.8 (i.e., a modest effective population size known with some, albeit limited, uncertainty). The environmental state for each population and generation was sampled from a standard normal distribution. I assumed a linear relationship between the environmental covariate and the selection differential, such that *S*_*jk*_ = *a* + *bx*_*jk*_ (i.e., Eqn. 1), and sampled values from the spike-and-slab priors for *a* and *b* for the simulated data sets using the same prior parameter values used for the ABC inference (lower and upper bounds of the slab component of the priors set to −0.1 and 0.1 and *π* = 0.5, see below).

I analyzed 2000 replicate simulations under these conditions (replicates differed in the values for selection, not in the trait genetic architecture or initial allele frequencies). Specifically, I generated 1.5 million simulations of evolution using the ABC simulation method, and treated the first 2000 as observed data. I applied the ABC inference procedure to each using the abc function from the R abc package (version 2.1) (Csillery *et al*., 2012). I used the local-linear regression method with a tolerance of 0.005, which retained 0.5% of the 1.5 million simulations (7500 samples) to form the posterior after regression adjustment. I summarized parameter estimates based on the posterior median (point estimate) and 95% equal-tail probability intervals (ETPIs). Performance of the method was then assessed by computing the mean absolute error (MAE), defined as the average absolute deviation between the true parameter value and its point estimate (posterior median), and the 95% interval coverage, that is the proportion of cases where the 95% ETPIs contained the true parameter value.

I further assessed the limitations of the method by analyzing four additional sets of simulations, each deviating in some way or ways from the standard conditions described above. One set considered the effect of enhanced genetic drift by replacing the re-scaled beta distribution above with one bounded by 100 and 200, and thus with an expected variance effective population size of *N*_*e*_ = 150. A second set assumed 1000 genetic loci might affect the focal trait, but postulated uncertainty in whether or not each had a causal effect. Specifically, each genetic marker was assigned a probability of association (i.e., *γ*_*i*_ or posterior inclusion probability) of 0.1. This results in an expectation of 100 causal variants but with uncertain identities. Third, I considered limited sampling, specifically five populations sampled across five generations. Fourth, I assumed a step selection function (Eqn. 3) rather than the linear function, with *π* = 0.5 and slab bounds of −0.05 and 0.05 for *c*. Each simulated data set (2000 per set of conditions) was analyzed as described above, with the exception that simulated genetic data with a step function for selection were analyzed with both the linear and step function models.

Lastly, I simulated additional data sets that deviated to greater extent from the assumed model. I specified the same genetic architecture, allele frequencies, effective population size and environmental data as for the standard conditions, but assumed that the environment defined an optimum for a Gaussian fitness function. Thus, instead of a simple selection differential (which only captures directional selection), selection would be a mixture of stabilizing and directional with specifics determined by the current composition of the population. The width of the Gaussian selection function (i.e., its standard deviation) was set to two or five (smaller standard deviations coincide with stronger stabilizing selection). The mean of the Gaussian function (i.e., that optimum phenotype) was specified by a linear model, *µ*_*jk*_ = *α* + *βx*_*jk*_. I considered two values for *α*, −0.762 or −0.562, and two values for *β*, 0 or 0.4. The first value for *α* (−0.762) coincides with the expected initial mean phenotype given the trait genetic architecture and allele frequencies, and thus with no (initial) directional selection in the average environment, whereas *α* = −0.562 should result in more instances of selection for larger trait values. Values for *β* were chosen to correspond with environment-independent (*β* = 0) and environment-dependent (*β* = 0.4) selection. I generated 20 simulated data sets for each combination of *α* and *β* (80 data sets total). This was done with an individual-based simulation written and implemented in R (version 3.5.1; code available from Dryad, DOI pending).

In these simulations, each generation genotypes for *N* = 1000 individuals were sampled from binomial probability distributions parameterized by the population allele frequencies. I then determined the phenotype of each individual by computing its polygenic score and adding environmental noise (to ensure the heritability of *∼*0.3) from a normal distribution. The relative fitness of each individual was then calculated from the Gaussian fitness function conditional on the environmental state. I then sampled *N*_*e*_ individuals, based on their relative fitness values, as survivors that determined the allele frequencies in the next generation. I repeated this procedure across 10 generation, 10 populations and 80 total simulations (20 replicates for each combination of selection function parameters). Because the values *α* and *β* for the selection function are for the phenotypic optimum (*µ*_*jk*_) rather than the selection differential (*S*_*jk*_), the method is not expected to estimate these values accurately. I thus instead compute approximate values for the true *a* and *b* from these simulations by fitting a linear model to the observed selection differentials from the simulations (i.e., from the change in the expected trait value from generation to generation), 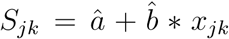 (the hat symbols denote that these are estimates of *a* and *b*, not parameter values defined by the model). I analyzed these 80 simulated data sets as described above, and measured performance by the ability to estimate â and 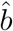.

I conclude my analysis of simulated data sets by illustrating how the proposed method can be used for model comparison. I chose (at random) one data set simulated based on the model with the linear selection function (and more generally, under the standard conditions). For this data set, I fit the linear selection function model (the true model) and the step selection function model (an alternative model), both as described above. I then generated 5000 samples from the posterior predictive distribution for the 10th to 90th percentiles of the distribution (over time and space) of change in polygenic scores based on each model. I compared these distributions graphically to the equivalent summary statistics computed from the observed data (in this case, the original simulated data set).

### Analysis of a seed beetle evolve-and-resequence experiment

I further tested the efficacy of the proposed method by using it to test for selection on female weight during an evolve-and-resequence experiment with the cowpea seed beetle, *Callosobruchus maculatus*. These insects infest stores of grain legumes (Tuda *et al*., 2014). Female beetles attach eggs to the surface of legume seeds. Hatching larvae burrow into the seed and must complete development in the single, natal seed.

My colleagues and I previously conducted an evolve-and-resequence experiment to analyze the evolutionary dynamics of host adaptation in *C. maculatus* when experimentally shifted to a stressful legume host, lentil (*Lens culinaris*), from an ancestral host, mung bean (*Vigna radiata*) (Rêgo *et al*., 2019). Low survival (*∼*1%) at the onset of the experiment caused a population bottleneck, but adaptive evolution quickly rescued the population from extinction with survival rates increasing to 90% by the F10 generation (Rêgo *et al*., 2019). Previous analyses of generation-scale allele frequency change from this experiment showed strong selection caused rapid adaptation and substantial evolutionary change at many genetic loci (Rêgo *et al*., 2019). Genome-wide association mapping in a back-cross (BC) mapping population derived from this line documented modest heritability for female weight at eclosion, with 17% of the trait variance explained by genetic markers (Rêgo *et al*., 2020). Female weight is one of several traits that often evolves during adaptation to lentil by *C. maculatus* (Messina *et al*., 2009; Messina & Jones, 2011; Messina & Durham, 2015). However, there was considerable uncertainty in individual genotype-weight associations, and we failed to detect excess overlap between SNPs associated with weight and those evolving most rapidly during the evolve-and-resequence experiment (Rêgo *et al*., 2020).

Here, I reanalyze these data with the proposed method, and specifically ask whether the method can detect selection on female weight and whether selection depends on population density (the hypothesized environmental covariate). Explicit data on population density are lacking, but qualitatively the population size had rebounded by the F4 generation, and thus I encode density as a binary covariate with a value of 0 for the first four generations and 1 thereafter. For this analysis, the data set included 10,409 SNPs sequenced in each of 10 samples (48 beetles per sample)–the founders (P), F1-F8 generations, and the F16 generation–and 251 sequenced female beetles that comprise the BC mapping population. Additional details regarding these data and experiments are provided by Rêgo *et al*. (2019) and Rêgo *et al*. (2020). SNP-genotype associations for the mapping population were estimated with the Bayesian sparse linear mixed model approach implemented in gemma (version 0.98) (Zhou *et al*., 2013). Consequently, for each SNP, there is an estimated posterior probability of association (i.e., posterior inclusion probability or PIP) and an estimated phenotypic effect conditional on the association. I assume the phenotypic variance in the experiment was equal to that in the mapping population (*s*^2^ = 0.43 mg; see the Discussion for limitations of this assumption). Bayesian allele frequency estimates were taken from Rêgo *et al*. (2020). I estimated the variance effective population size over the course of the experiment based on patterns of allele frequency change between the P and F16 generation using the Bayesian approach implemented in varNe (version 0.9) (Gompert & Messina, 2016).

I used the proposed method to estimate selection on female weight. The model was fit in two ways, using either (i) model-averaged effect estimates (i.e., taking the product of the posterior inclusion probability and effect estimate as a certain, model-averaged effect estimate) or (ii) integrating over uncertainty in genotype-phenotype associations. My main focus was on the analysis with population density (a qualitative, binary indicator variable) as an environmental covariate, but I also considered a null model with directional selection only and a model with a random environmental covariate (with environmental states drawn from a standard normal distribution). I bounded the priors on the selection model parameters (*a* and *b*) by −2.0 and 2.0 (here *S* is in mg). In each case, I based the analysis on one million simulations. In the models with uncertainty in genotype-phenotype associations, observed summary statistics were obtained for 100 independent samples of genotype-phenotype associations (based on their PIPs). Inferences were based on 1000 samples from the posterior after apply the local-linear regression adjustment in abc (Beaumont *et al*., 2002; Csillery *et al*., 2012).

## Results

### Results from stimulated data

The core simulations (i.e., those based on the model) resulted in weak to modest phenotypic selection, such that the average, absolute intensity of selection was less than 0.01 for 55% of the simulations and had a maximum value of *∼*0.1 (Fig. S1). Variability in selection across space and time was of the same order of magnitude as the average, absolute intensity of selection, consistent with Siepielski *et al*. (2009) and Siepielski *et al*. (2013) (mean standard deviation in |*S*_*jk*_| = 0.025, maximum = 0.10) (Fig. S1). The average selection intensity on individual genetic loci (measured by |*s*_*ijk*_|) was about an order of magnitude lower, and rarely exceeded 0.01 (Fig. S2).

Under the standard conditions (i.e., linear selection function, 10 populations, 10 generations, 100 causal loci, 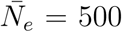, *h*^2^ = 0.3), I obtained precise and accurate estimates of the mean selection differential (*a*) and environmental effect (*b*) (Figs. 2). For example, the MAE (i.e., the average absolute deviation between the true and estimated parameter value) was 0.0028 for *a* and 0.0025 for *b*. Moreover, the 95% interval coverage was 0.95 and 0.96 for *a* and *b*, respectively. High accuracy was mostly independent of the strength of these effects (Fig. S3). Estimates of the average selection on genetic loci were similarly accurate with a MAE of 0.00031 and 95% interval coverage of 0.96 for the standard conditions (Fig. 3). There was a slight decrease in the accuracy of the estimates for simulations with reduced *N*_*e*_ (i.e., cases where drift was more important), uncertainty in genetic architecture, reduced spatial and temporal sampling or with the step selection function (Fig. 4). For example, MAE increased, approximately doubling, with reduced sampling (MAE = 0.0064 for *a* and 0.0061 for *b*), though 95% interval coverage remained high (*∼*95%).

**Figure 2:**
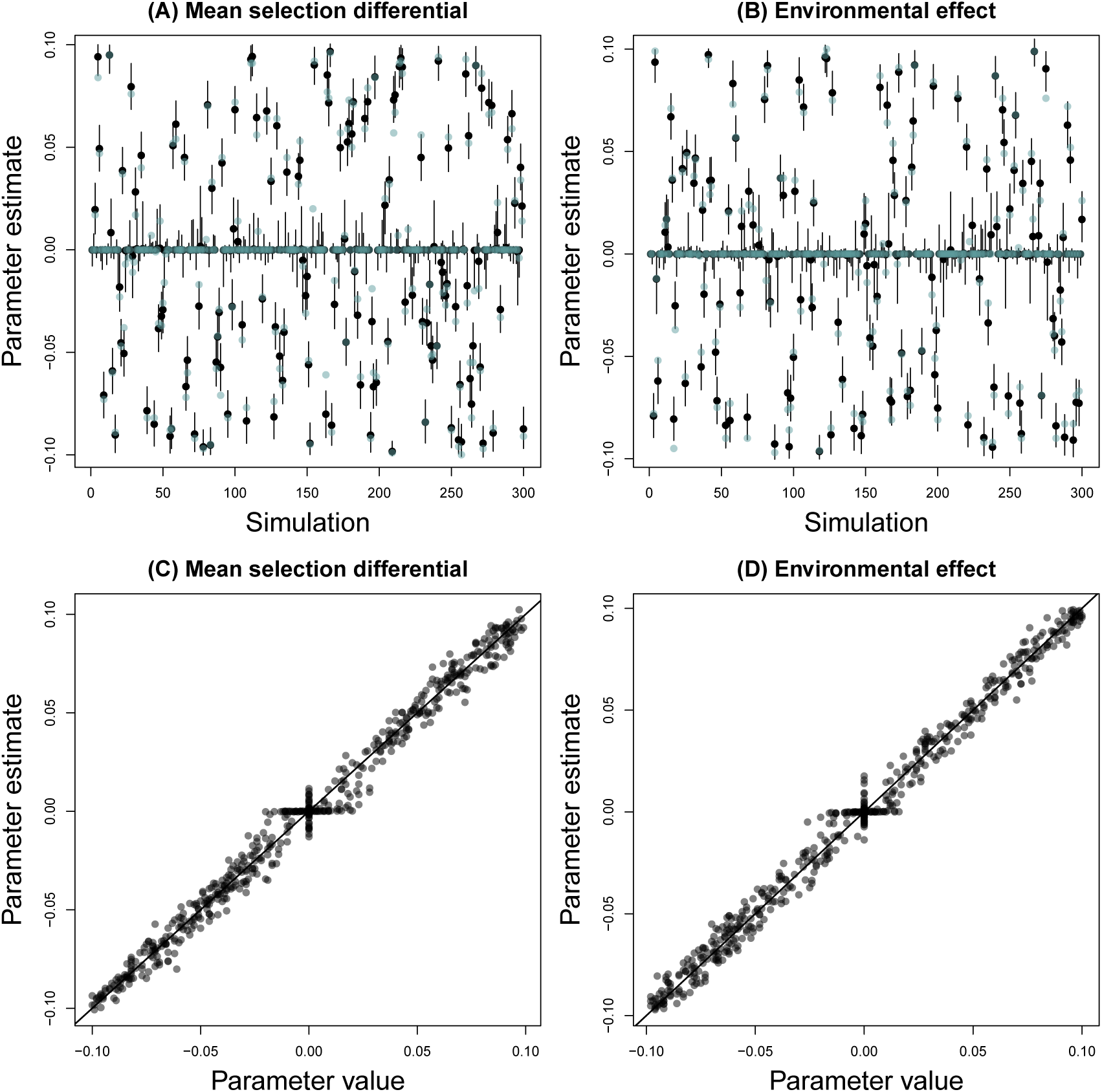
Plots summarize performance of the method under the standard simulation conditions. Panels (A) and (B) show the true (blue points) and estimated (black points with vertical lines for 95% equal-tail probability intervals [ETPIs]) parameter values for the selection coefficients *a* and *b*, respectively. Results are shown for 300 representative simulations. Panels (C) and (D) provide scatterplots depicting the relationship between the true parameter values (x-axis) and the point estimate (posterior median, y-axis) for *a* (C) and *b* (D). A one-to-one line is shown. Parameter estimates were highly correlated with their true values (Pearson *r >* 0.99).

**Figure 3:**
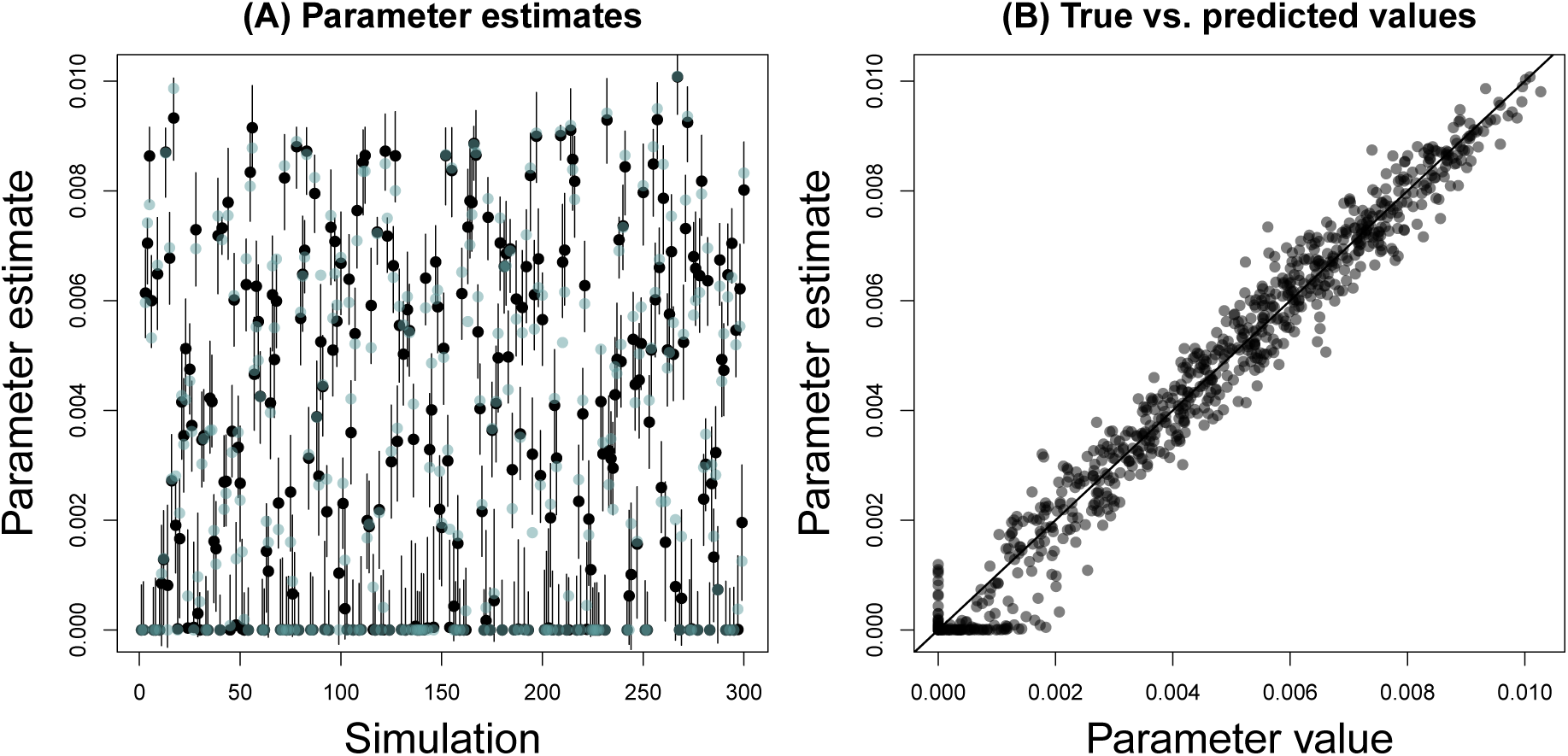
Plots summarize performance of the method in terms of selection on genetic loci under the standard simulation conditions. Panel (A) shows the true (blue points) and estimated (black points with vertical lines for 95% equal-tail probability intervals [ETPIs]) mean absolute intensity of selection on the genetic loci. Results are shown for 300 representative simulations. Panel B provides a scatterplot depicting the relationship between the true mean intensity of selection on genetic loci (x-axis) and the point estimate (posterior median, y-axis). A one-to-one line is shown. Parameter estimates were highly correlated with their true values (Pearson *r* = 0.99).

**Figure 4:**
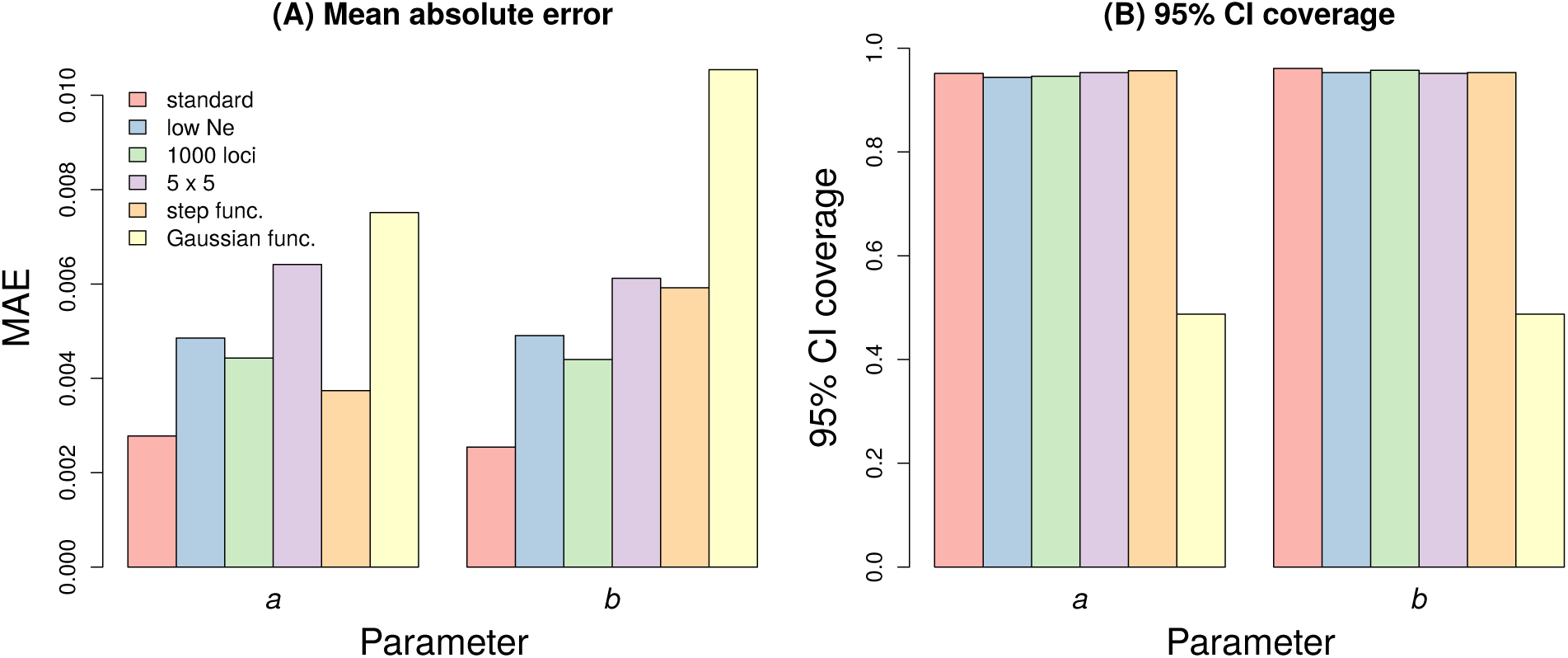
Barplots depict how performance of the method depends on simulation conditions. Results are shown for mean absolute error (MAE) (A) and 95% interval coverage (B).

There was a greater reduction in performance for simulations based on the Gaussian fitness function, that is where the analysis model differed from the generative evolutionary process used to simulate the data (Figs. 4 and 5). In particular, MAE increased to 0.0075 for *a* and 0.0105 for *b*, and the 95% interval coverage dropped to *∼*0.49 for both. Despite this drop in performance, the parameter estimates were still indicative of the “true” parameter values (recall that â and 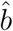 are not true parameter values in the same sense for these data sets, but rather best approximations when forcing a linear model for the selection differential). For example, Pearson correlations between true and estimated values (posterior medians) for *a* and *b* across the 80 data sets simulated under this model were 0.80 (95% confidence interval = 0.71–0.87) and 0.97 (95% confidence interval = 0.96–0.98), respectively.

**Figure 5:**
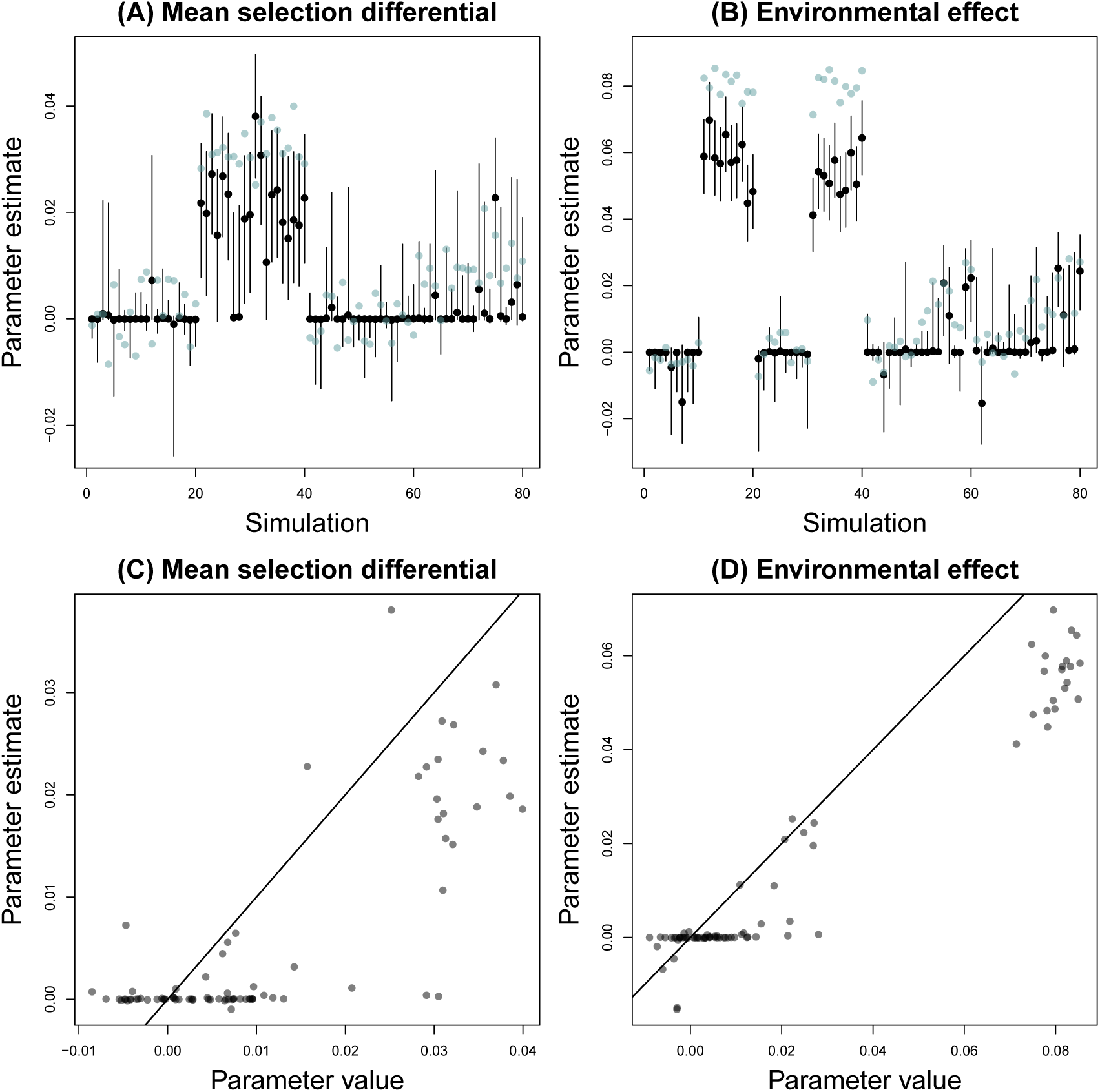
Plots summarize performance of the method for data simulated assuming a Gaussian fitness function with an optimum determined by the environment. Panels (A) and (B) show the true (blue points) and estimated (black points with vertical lines for 95% equal-tail probability intervals [ETPIs]) parameter values for the selection coefficients *a* and *b*, respectively. Panels (C) and (D) provide scatterplots depicting the relationship between the true par meter values (x-axis) and the point estimate (posterior median, y-axis) for *a* (C) and *b* (D). A one-to-one line is shown.

A comparison of the posterior predictive distribution for models with either linear or step selection functions fit to a simulated data set where the true function was linear illustrates how the proposed method can be used for model comparison (see Fig. S4 for the model fits). As expected, the true summary statistics fell closer to the center of the posterior predictive distribution for the true linear model than the step function model (Fig. 6). This held across the various percentiles considered for the summary statistics.

**Figure 6:**
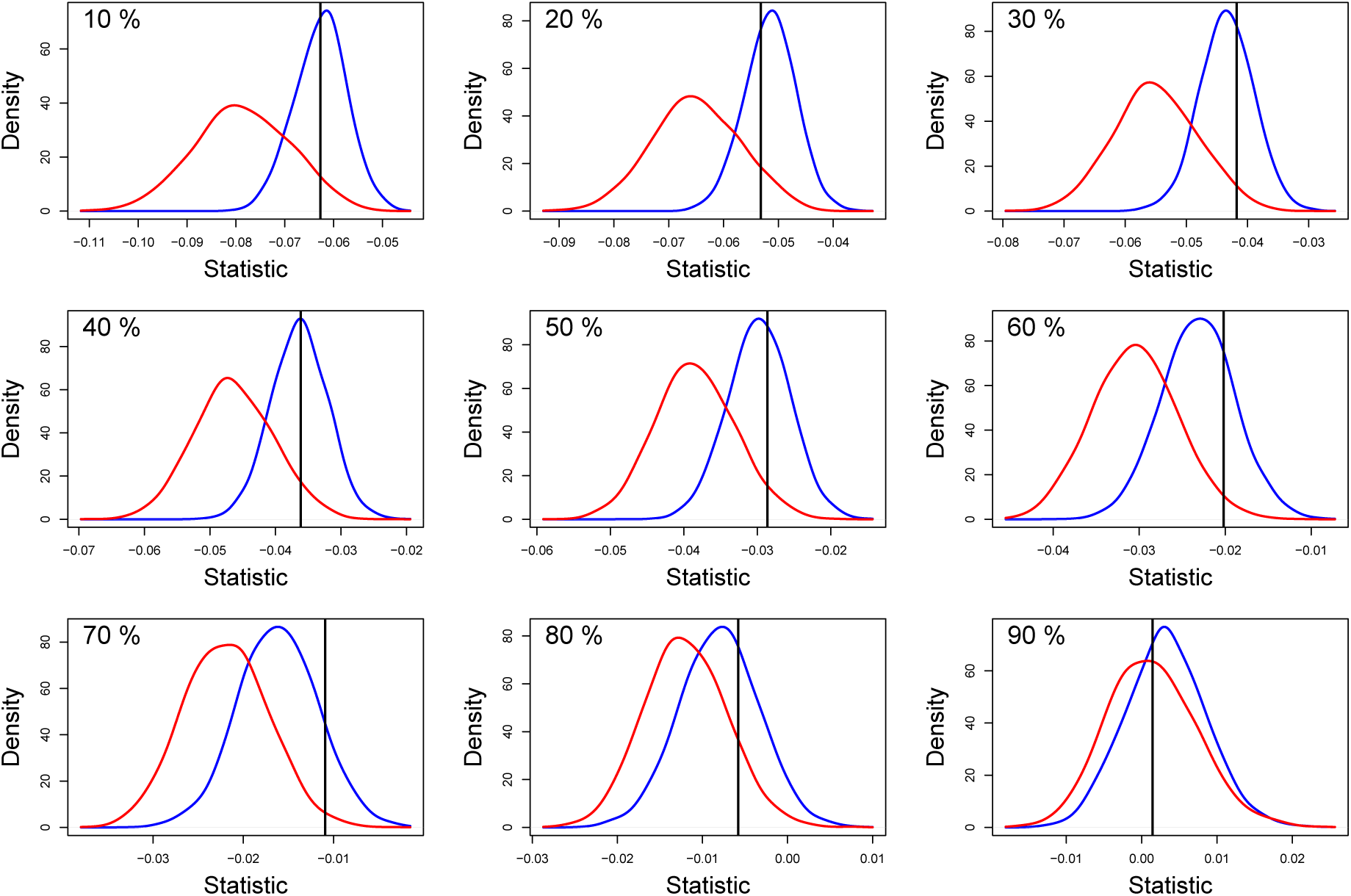
Graphical summary of a posterior predictive check. In each panel, the posterior predictive distribution is shown for the linear selection model (blue) and the step selection model (red). The former is the true model. The summary statistic from the data set used to fit the models (i.e., the observed data) is shown as a vertical black line. Each plot shows results for a different summary statistic, specifically a different percentile of the distribution of change in the mean polygenic score across populations and generations (the percentile is given in the upper left corner of each plot).

## Results from the seed beetle evolve-and-resequence experiment

I found compelling evidence of strong selection for increased female weight when using model-averaged genotype-phenotype effects (Fig. 7A). Estimates of the average selection on weight were mostly insensitive to my choice of an environmental covariate, with estimate of *a* equal to 1.95 (95% ETPI = 1.70–2.01), 1.99 (1.85–2.03) or 2.00 (1.88–2.06) when using population density, a random covariate or no covariate, respecitvely. In contrast, fully accounting for uncertainty in genotype–phenotype associations dramatically reduced the evidence for selection (Fig. 7A). When fully accounting for uncertainty in the genotype-phenotype map in simulations and the observed data, the estimate of *a* was *∼*0 (posterior mean = 0.156, 95% ETPI = −0.69–1.57, posterior prob. *a >* 0 = 0.62). Using the mean of summary statistics across replicate genotye-phenotype maps for the observed data decreased uncertainty in *a* some and increased the evidence for *a >* 0 slightly, but had little effect on the point estimate, 0.0003 (posterior mean = 0.020, 95% ETPI = −0.00–0.28, posterior prob. *a >* 0 = 0.83). This discrepancy between results based on model-averaged phenotypic effects and fully accounting for uncertainty in the genotype-phenotype associations is consistent with calculations of mean breeding values or polygenic scores over the course of the experiment and likely reflects the great uncertainty in gentoype-phenotype associations in this data set (the highest SNP posterior inclusion probability was 0.03 and only 13 SNPs had posterior inclusion probabilities *>* 0.01) (Fig. S5).

**Figure 7:**
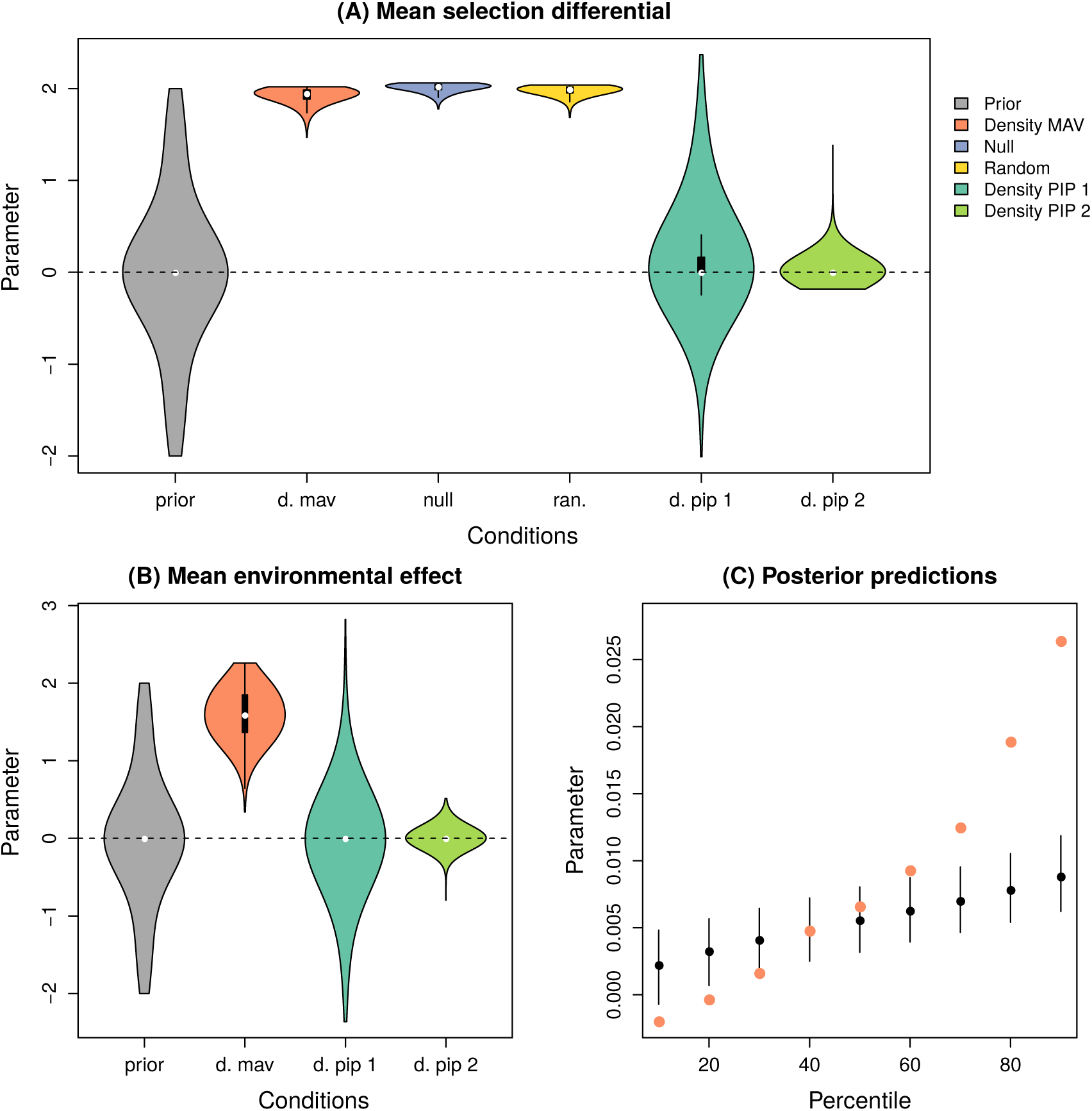
Plots summarize the application of the proposed method to the *C. maculatus* data set. Violin plots in panel (A) depict prior (gray) or posterior (other colors) distributions for the mean selection differential parameter, *a*. Posteriors are shown with density as a covariate and with model-averaging (Density MAV) or uncertainty with the distribution (Density PIP 1) or mean (Density PIP 2) observed summary statistics, and for no covariate (Null) or a random covariate (Random). White dots denote the median, black boxes denote the 1st and 3rd quartile, vertical bars extend to the minimum and maximum value or 1.5× the interquartile range, and kernel densities show the full prior or posterior distribution. Similar plots in panel (B) show the prior and posteriors for the environmental effect, *b*. Panel C summarizes the posterior predictive check. Colored points shown percentiles of change in the mean polygenic score for the observed data (using model-averaging), and black dots and lines show the median and 95% intervals from posterior predictive simulations.

Similar to the results for *a*, I found evidence that selection varied by population density when using model-averaged effects, but not when accounting for uncertainty in genotype-phenotype associations (Fig. 7B). Even when using model-averaged effects, I found no evidence that selection on weight varied as a function of the random covariate (*b* = −0.19, 95% ETPI = −1.33–1.31, posterior prob. *b <* 0 = 0.69). Regardless of the specific model, estimates of the average and variability of selection on individual loci were always low (*<* 0.001). Lastly, a posterior predictive check based on the best performing model, that is the population density model with model-averaged effects, underestimated the variability in change in the expected polygenic score across generations (Fig. 7C).

## Discussion

Rapid evolution on ecological time scales is at least somewhat common (Endler, 1986; Thompson, 2013; Bergland *et al*., 2014; Grant & Grant, 2014; Hendry, 2016; Nosil *et al*., 2018), but remains understudied from a population genomic perspective (Messer *et al*., 2016). This is perhaps not surprising, as most existing methods were designed to detect weaker selection operating in a consistent manner over longer periods of time. I proposed and evaluated a statistical method to help fill this gap. Critically, the method connects environmental drivers of selection with traits and patterns of evolutionary change in a population genomic context. Taking a population genomic approach has the advantage of estimating selection on genetic variants (not just traits) and of mostly disentangling temporal variation in traits due to evolutionary change versus plasticity. Via simulations, I showed that the proposed method can generate very accurate and precise estimates of fluctuating selection on polygenic traits. However, performance depended on simulation conditions and is generally expected to be sensitive to details of the data input. I discuss performance at length below, along with current limitations and possible extensions of the proposed method.

### Overall performance

The proposed method was computationally efficient, and I do not foresee problems arising even when applying it to large, whole-genome data sets. As an example, I was able to generate one million ABC simulations for the *C. maculatus* data set in *∼*300 CPU minutes on a standard desktop computer (Intel Core i7-3930K CPU at 3.20 GHZ *×*12 with 64 GB RAM and running Ubuntu 18.04.4 LTS). Moreover, each simulation is independent, and thus massive parallelization of the ABC simulations is trivial. Computational time should scale linearly with the number of populations and generations, and with the number of genetic variants associated with or affecting the phenotype. Notably, the number of phenotype-associated genetic markers will likely be on the order of hundreds to perhaps tens of thousands even when whole-genome sequences are analyzed (i.e., not every genetic variant affects a trait).

In terms of accuracy of inference, the method performed best under the standard simulation conditions (i.e., 10*×*10 populations*×*generations, moderately large *N*_*E*_, and 100 known causal variants). I documented minor degradation of performance with more limited space-time sampling (5*×*5 populations*×*generations) or reduced effective population size (i.e., increased genetic drift). Deviations between the assumed and true selection function and uncertainty in genetic architecture had greater effects, and I discuss each in turn next.

### Performance and the selection function

Performance of the method declined substantially when applied to simulations of environment-dependent stabilizing selection described by a Gaussian fitness function. For example, the true parameter value (or at least the linear approximation to the true parameter value) was not within the 95% ETPIs of the estimate about half of the time. Nonetheless, the estimates tended to be correct qualitatively, that is, the direction and approximate magnitude of the mean selection differential and environmental effect were correctly estimated. Future work could allow for additional selection functions, but even then, quantitative estimates of selection will only be strictly interpretable when the function assumed matches (or very closely approximates) reality. Estimates of selection function parameters cannot be combined across functions as these parameters do not have a consistent meaning, but Bayesian-model averaging for derived parameters is possible.

Perhaps even more important from a practical perspective, the estimates of selection, or at least the environment-dependent component of selection, depend on correctly identifying the environmental covariate responsible for fluctuations in selection. In some cases, such as negative or positive frequency-dependent selection, this might be relatively easy. And a recent meta-analysis suggests that some aspects of the environment, most notably precipitation, might generally be important drivers of variation in selection (Siepielski *et al*., 2017). Nonetheless, it will often be difficult to designate a single environmental covariate *a priori*. Alternative models with different covariates could be fit and evaluated based on their posterior predictive distributions. Indeed, the relatively poor fit of the population density model to the *C. maculatus* data set was uncovered based on the posterior predictive distribution (unfortunately, I lack a clear, alternative covariate to explain variability in change in the mean polygenic score over time in this data set). However, caution is warranted as considering many possible environmental covariates may uncover spurious relationships. When many covariates are tested, it would be prudent to divide the data into training and validation sets for cross-validation. Lastly, in some cases variation in selection likely results from the interaction of multiple environmental factors (e.g., Benkman & Siepielski, 2004; Benkman *et al*., 2010; Nosil *et al*., 2018). I intend to extend the method and `fsabc` software to allow for multiple covariates and their interactions in the future.

### Performance and the genotype-phenotype map

Uncertainty in trait genetic architecture had a modest effect on performance of the method with simulated data, but had a profound effect on the analysis of the *C. maculatus* experiment. At least two factors likely contributed to this discrepancy. First, the simulations involved 1000 genetic markers each having a modest probability of being associated with the trait (all PIPs = 0.1), whereas more than 10,000 SNPs had small but non-zero probabilities of association with female weight in *C. maculatus* (minimum = 0.001, maximum = 0.030, median = 0.004). Thus, the *C. maculatus* data exhibited much greater uncertainty in the genotype-phenotype map than the simulated data. Second, the method assumes the posterior inclusion probabilities (PIPs) across genetic markers are independent (i.e., all information arises from the marginal probabilities of association). This assumption was valid for the simulated data, but was likely violated for *C. maculatus* data set. With the Bayesian sparse linear mixed model approach used to estimate genotype-phenotype associations for *C. maculatus* weight, sets of SNPs in the model (i.e., with non-zero effects) in any given Markov chain Monte Carlo (MCMC) step are unlikely to be associated with the same causal variant (or in high LD with each other) (Zhou *et al*., 2013). Instead, on average each causal variant should be represented by *∼*1 SNP marker in any given MCMC step. Information on sets of SNPs (not just individual SNPs) is not encoded in the PIPs. Thus, the marginal association probabilities jettison information from the full, joint posterior and sampling sets of SNPs for the proposed method based on these could result in poor inference in cases where the marginal posteriors are misleading.

I see several ways to overcome the problem of uncertainty in the genotype-phenotype map. First, additional experiments could be conducted, including genetic manipulations or fine-scale genetic mapping, to refine an initial set of candidate genetic markers or QTL regions (e.g., Linnen *et al*., 2013; Barrett *et al*., 2019). Second, when sets of linked SNPs are associated with a trait, all but the most strongly associated could be dropped and the sum of the posterior inclusion probabilities across the set could be assigned to the retained SNP (this could be repeated for different sets of traits). This should work best when genetic markers fall into multiple, easy to delineate, sets of trait-associated markers (which was not the case for the *C. maculatus* data set; Rêgo *et al*., 2020). Third, the method could be modified to work with samples from the joint posterior for genotype-phenotype associations. Such information can be extracted from the MCMC output of common polygenic GWA models (including gemma; Zhou *et al*., 2013). I plan to implement this final approach in a future version of `fsabc`. Short of this, my results suggest that using model-averaged effect estimates is likely the better solution when dealing with considerable uncertainty in the genotype-phenotype map. At minimum, this approach allowed us to detect selection in *C. maculatus*.

An additional key consideration of the genotype-phenotype map concerns the choice of a mapping population. I expect the proposed method to perform best when the mapping population harbors the same genetic variants segregating at the same frequencies, and the same phenotypic variance, as the populations included in the population-genomic time series. Thus, mapping in those actual populations will often be the best choice. Indeed, differences between the genetic composition of the *C. maculatus* mapping population (a BC between the source population and the lentil-adapted line) and the focal line during the course of the experiment likely further limits quantitative interpretation of the magnitude of selection on weight during the experiment (more generally, mapping populations generated from crosses might often violate the linkage and Hardy-Weinberg equilibrium assumptions for equating average effects with average phenotypic excess). Of course, if the genetic composition of the populations changes considerably over the course of the study (perhaps because of sustained directional selection) or if the phenotype is highly plastic, a single genotype-phenotype map may be limiting regardless of the mapping populations (e.g., Coop, 2019). In such cases, it should be possible to include multiple genotype-phenotype maps (for different populations, generations, or environments). Doing so would allow selection and the genotype-phenotype map to depend on the environment and would thus facilitate the analysis of plastic traits. I hope to include this in a future version of `fsabc`.

### Conclusions

Combining population-genomic time-series data with data on trait genetics and the environment has the potential to advance our understanding of how selection acts in nature and drives evolution across the genome. I hope that the proposed method contributes to this aim. I expect the biggest limitation at present for the use of this (or related) method is a lack of appropriate data. Large-scale spatio-temporal population genomic data sets are still uncommon, especially for natural populations. However, advances in the analysis of ancient or museum DNA and the expanding field of evolve-and-resequence experiments may help overcome this limitation (e.g., Bi *et al*., 2013; Barghi *et al*., 2017; Sproul & Maddison, 2017; Cridland *et al*., 2018; Rêgo *et al*., 2019). Moreover, few systems currently combine such sampling with detailed studies of trait genetics and the environment. Nonetheless, I hope that methods, like the one proposed here, help illustrate the potential value of such combined data sets, and more generally of long-term studies in the age of genomics (see e.g., Grant & Grant, 2014).

## Acknowledgments

This manuscript was improved by comments from A. Bergland and C. Nice. This research was funded by the National Science Foundation (DEB-1638768 and DEB-1844941 to ZG). The support and resources from the Center for High Performance Computing at the University of Utah are also gratefully acknowledged.

## Data Accessibility

The `fsabc` software, manual and an example data set are available from GitHub (https://github.com/zgompert/fsabc.git). Additional scripts and simulated data sets analyzed in this manuscript are available from Dryad (DOI pending).

## Author Contributions

ZG designed the study, developed the proposed method, wrote the computer software, generated and analyzed the simulated data, analyzed the *C. maculatus* data, and wrote and revised the manuscript.

## Supplemental material for

### Supplemental Tables and Figures

**Figure S1:**
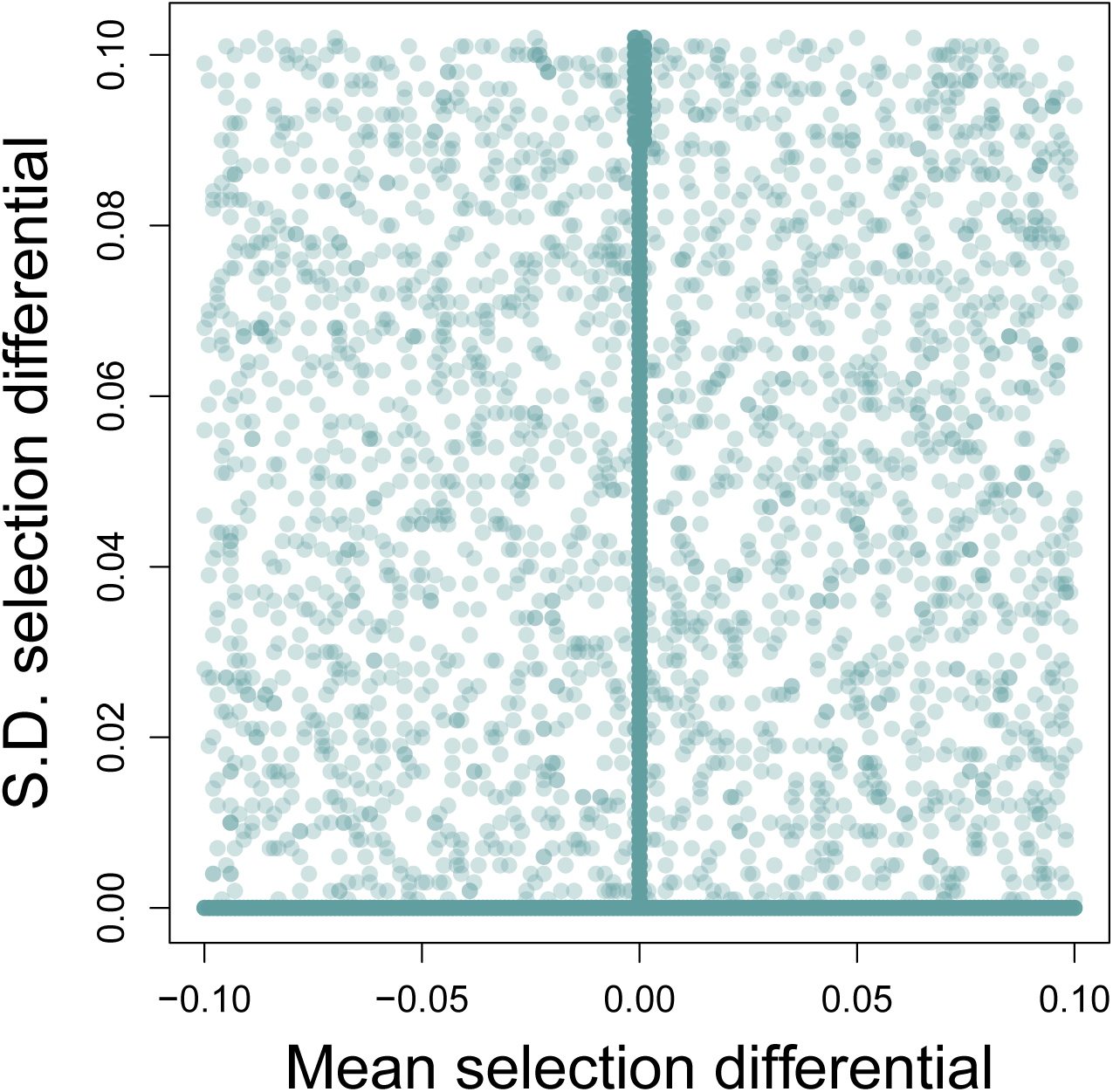
The scatterplot shows the mean absolute selection differential versus the standard deviation in the absolute selection differential for a random sample of 10,000 simulations. The high density of points near 0 on the x- and y-axes reflects the spike component of the spike-and-slab prior.

**Figure S2:**
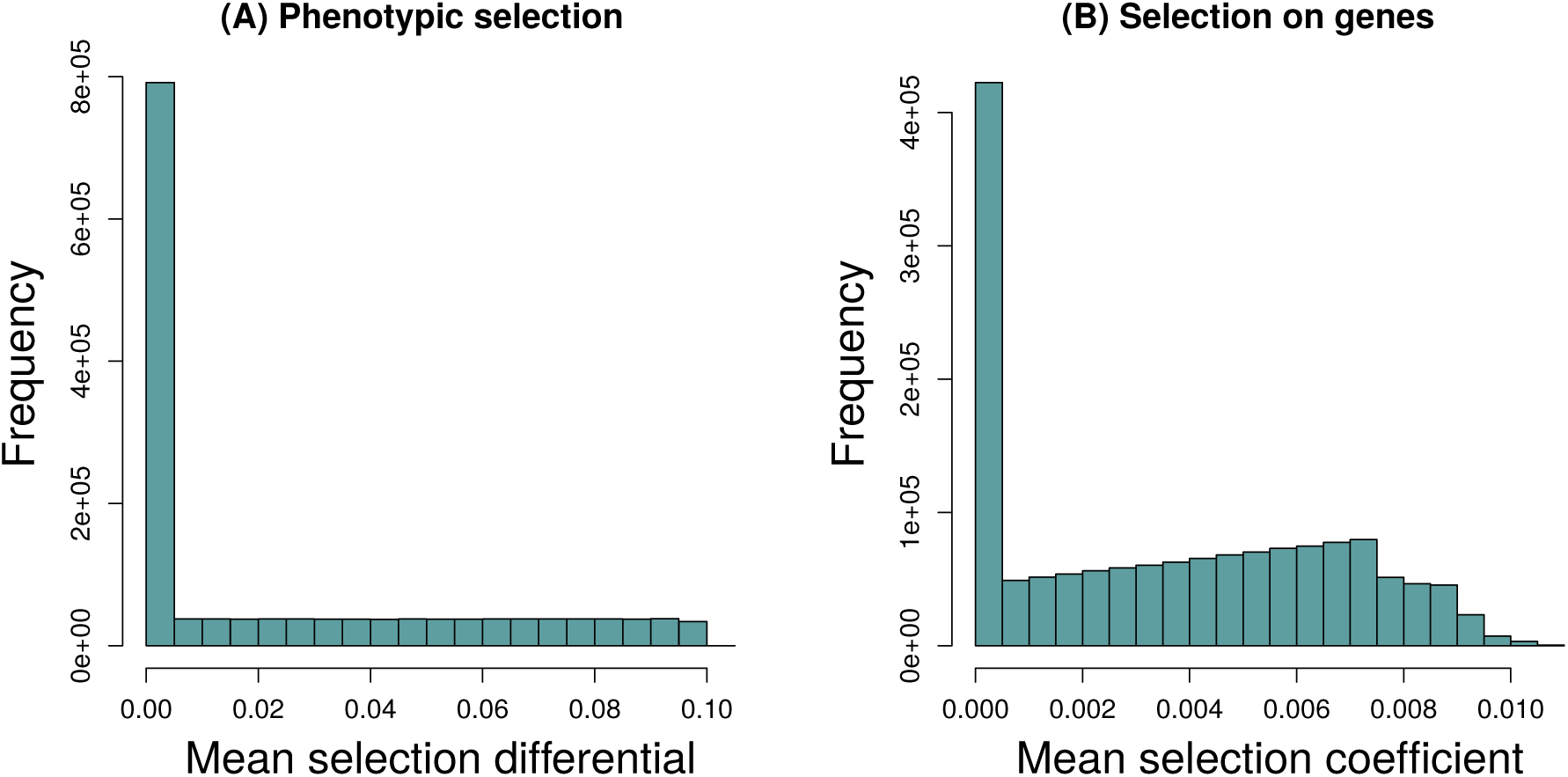
Histograms compare the distribution of the average intensity of selection on the focal trait (as measured by |*S*_*jk*_|) (A) and the average intensity of selection on underlying genetic loci (as measured by |*s*_*ijk*_|) (B).

**Figure S3:**
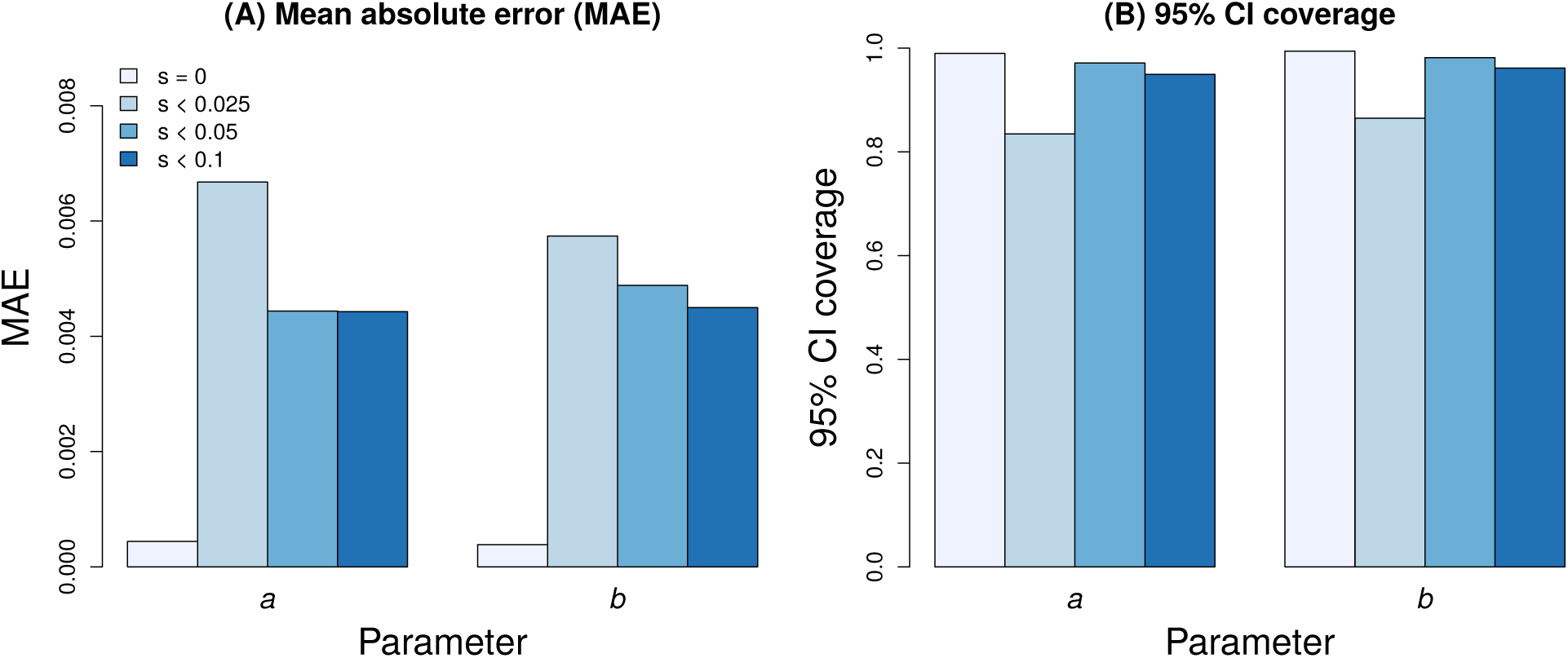
Plots show the effect of the intensity of selection (measured by *a* or *b*) on the performance of the method. Here, I stratify simulations based on *a* or *b* = 0, or falling into one of the following intervals, *>*0– ≤0.025, *>*0.025– ≤0.05, *>*0.05– ≤0.1 (see legend where I use *s* as generic notation for *a* or *b*). Results are shown for mean absolute error (MAE) (A) and 95% interval coverage (B).

**Figure S4:**
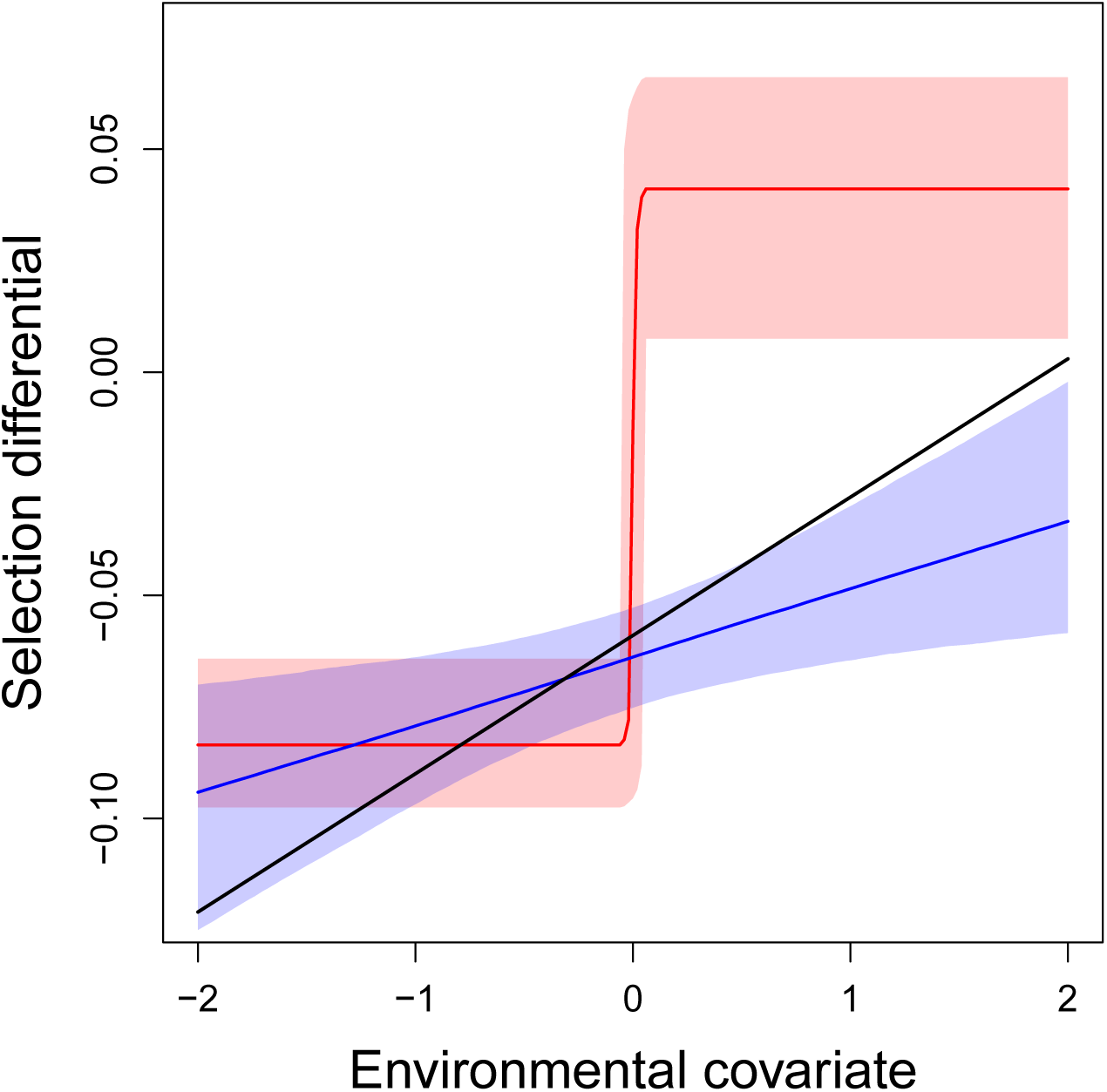
Posterior estimates of the selection function under alternative models. Colored lines (posterior median) and shaded regions (95% equal-tail probability intervals) show estimates of the linear (blue) and step (red) functions for the selection differential. The true model used to simulate the data had a linear selection function, which is shown as a black line.

**Figure S5:**
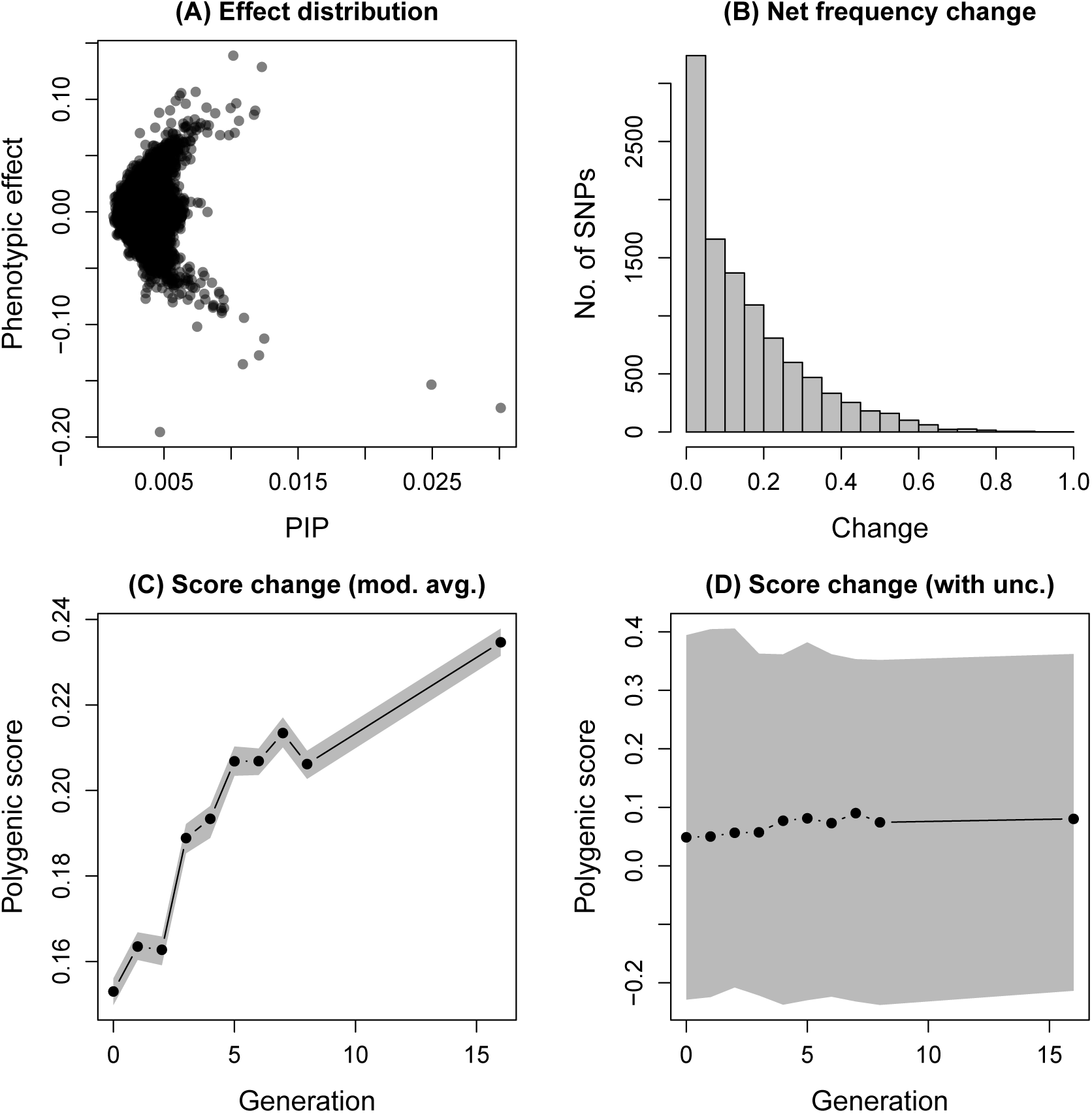
Summary of trait (female weight) genetics and evolutionary change in *C. maculatus*. Panel (A) shows the posterior inclusions probability (PIP) for each of 10409 SNPs in the genotype-phenotype model for weight versus the estimated effect of each SNP on weight. Panel (B) shows the distribution, across the 10409 SNPs, of net allele frequency change from the founders of the lentil line to the F16 generation. Panels (C) and (D) show the observed change of the mean polygenic score relative to the mean of the mapping population in the experimental population. Dots denote point estimates and the shaded gray regions denote 90% credible intervals. Results in (C) used model-averaged effect estimates and uncertainty arises only from uncertainty in allele frequencies. In contrast, results in (D) are based on repeat sampling of sets of actually associated SNPs based on their PIPs and thus fully account for uncertainty in the genotype-phenotype map.

